# T cell protrusions enable fast, localised initiation of CAR signalling

**DOI:** 10.1101/2025.07.08.662959

**Authors:** Carmen Rodilla-Ramirez, Giorgia Carai, Eleanor Fox, Amin Zehtabian, Helen Adam, Katja Dallio, Helge Ewers, Xiaolei Su, Francesca Bottanelli

## Abstract

Actin-rich protrusions densely cover the surface of T cells and are well characterised for their role in cell migration. However, recent studies have uncovered their role in antigen surveillance and immune signalling initiation. To investigate how membrane protrusions initiate and contribute to signalling, from the first cell-cell contact to immunological synapse formation, we performed dynamic imaging experiments of endogenously tagged signalling proteins in T cells. To quantitatively capture the early dynamics of cell-cell interactions, we employed HER2-CAR-expressing T cells targeting HER2^+^breast cancer cells. By harnessing live-cell imaging and super-resolution stimulated emission depletion (STED) microscopy we were able to capture topological membrane changes and their correlation with mesoscale protein rearrangements over time. Our findings indicate that, prior to activation, key molecular players in T cell activation, including the kinase Lck, the phosphatase CD45 and the adaptor LAT, as well as the exogenously expressed CAR, lack any enrichment in actin-rich protrusions. However, upon initial contact of the T cell with the target cell, a dynamic and fast rearrangement of the surface receptors, phosphatases, and kinases occurs within the protrusions, ensuring a rapid and effective initiation of the immune signalling cascade. The rapid clustering of the HER2-CAR occurs preferentially within protrusions rather than flat membrane regions and is accompanied by enhanced recruitment of the kinase ZAP-70 and LAT. While the localisation of the kinase Lck remained unchanged, protrusion-cell contacts trigger a pronounced exclusion of the phosphatase CD45, an effect observed both with and without the cytosolic signalling domain of the CAR. Overall, the signalling machinery rearranged more rapidly and efficiently at contacts mediated by protrusive structures compared to non-protrusive regions. Together, our data provide a quantitative framework illustrating how signalling proteins are dynamically reorganised to facilitate CAR-mediated activation within these specialised structures.

## Introduction

T lymphocytes play a central role in the adaptive immune response, using their T cell receptor (TCR) to recognise foreign antigen peptides presented by major histocompatibility complexes (MHCs) on the surface of antigen-presenting cells (APCs). Upon sustained TCR interaction with peptide MHC (pMHC), the Src family kinase Lck phosphorylates immune receptor tyrosine-based activation motifs (ITAMs) on the cytoplasmic domains of the TCR, creating docking sites for the kinase ZAP-70 (Iwashima et al., 1994). Once at the TCR, ZAP-70 is activated by Lck and subsequently phosphorylates substrates such as the linker for activation of T cells (LAT) (Zhang et al., 1998). LAT then serves as a scaffold to drive biomolecular condensation (Su et al., 2016) from which downstream TCR signalling pathways originate (Balagopalan et al., 2013; Balagopalan et al., 2015). Beyond the molecular components of T cell signalling, T cell function is also influenced by its complex membrane topography. The T cell surface is covered by a heterogenous set of actin-rich protrusions (Alexander et al., 1976; Kenney et al., 1986; Majstoravich et al., 2004) that change in density and/or morphology in response to the physical and chemical context (Brown et al., 2003; Real et al., 2007; Shulman et al., 2009), as well as the activation status of the cell (Fritzsche et al., 2017; Sanderson and Glauert, 1979). Protrusions have traditionally been studied in the context of T cell migration through blood and tissues (Carman et al., 2007; Caswell and Zech, 2018; Dupré et al., 2015; Song et al., 2014). However, studies over the past decade have highlighted a crucial role of actin-rich protrusions in immune signalling {Stinchcombe, 2023 #8}(Aramesh et al., 2021; Beppler et al., 2023; Cai et al., 2022; Cai et al., 2017; Jenkins et al., 2023; Kim et al., 2018; Orbach and Su, 2020; Sage et al., 2012; Tamzalit et al., 2019). Actin-rich protrusions, commonly referred to as T cell microvilli, have been shown to rapidly scan the APC surfaces and stabilise upon TCR-pMHC binding (Cai et al., 2017). Furthermore, they are thought to be the preferred energetic solution for penetrating the glycocalyx of both T-cell and target cell (Göhring et al., 2022; Pettmann et al., 2018), which can physically hinder intercellular receptor-ligand interactions (Ardman et al., 1992; Bell et al., 1984). Protrusions have been shown to “punch” into the target cell and bring T cell and target membranes closer to allow surface receptor interactions (Sage et al., 2012; Sanderson and Glauert, 1979). Disruption of the T cell actin cytoskeleton delays this close contact formation (Jenkins et al., 2023) suggesting the need for actin-driven forces for cell-cell interactions. An abundant component of the T cell glycocalyx is the large phosphatase CD45. Size-dependent exclusion of CD45 from TCR-pMHC interaction sites, shown at different timepoints of T cell activation (Chang et al., 2016b; Razvag et al., 2018; Varma et al., 2006), has been proposed to be the driving element for TCR triggering, according to the kinetic segregation model (Davis and van der Merwe, 2006).

A quantitative dynamic framework describing macromolecular rearrangements in relation to membrane topography on the surface of T cells, from the first contact to immunological synapse formation and in an unperturbed living cell, is currently lacking. The localisation of signalling proteins has conventionally been imaged by overexpressing proteins of interest fused to a fluorescent tag (Gudipati et al., 2020; Razvag et al., 2018) or via immunostaining of fixed cells (Beppler et al., 2023; Dong et al., 2020; Ghosh et al., 2020; Jung et al., 2016a; Jung et al., 2021). While fixation has been instrumental for high-resolution imaging, it may alter membrane structures (Ichikawa et al., 2022; Schnell et al., 2012; Wong-Dilworth et al., 2023) underscoring the importance of complementary live-cell techniques. Single molecule localisation microscopy (SMLM) has been broadly employed to assess the nanoscale localisation and clustering behaviour of signalling molecules on the plasma membrane of T cells, offering powerful insights (Lillemeier et al., 2010; Sherman et al., 2011; Williamson et al., 2011)—though careful controls are needed to account for potential clustering artefacts (Rossboth et al., 2018).

The localisation of signalling molecules in T cell protrusions prior to activation has been an active area of investigation, due to the possibility that T cells may enrich signalling molecules at their tips to facilitate initial antigen sensing. While some studies report enrichment of TCR and pre-exclusion of CD45 in protrusions in resting T cells (Ghosh et al., 2020; Jung et al., 2016b; Jung et al., 2021), others present contrasting findings (Cai et al., 2022), reflecting the complexity and dynamic nature of these structures. Furthermore, most studies on actin-rich protrusions have relied on imaging T cell interactions with coated glass surfaces (Chang et al., 2016b; Razvag et al., 2018) or lipid bilayers (Beppler et al., 2023; Cai et al., 2022; Cai et al., 2017; Jenkins et al., 2023), where membrane topology in relationship to domains enrichment is hard to infer due to the lack of axial resolution in most conventional and super-resolution imaging methods. While this system has greatly advanced our understanding of T cell signalling, it highlights the need for approaches that capture the complex topography of the T cell plasma membrane during more physiological cell-cell interactions.

Here, we employed CRISPR-Cas9 to endogenously tag key signalling proteins such as Lck, ZAP-70, LAT or CD45, and performed quantitative live-cell confocal and super resolution [stimulated emission depletion (STED)] microscopy to develop a quantitative model of their dynamics and nanoscale redistribution during Human Epidermal Receptor 2 (HER2)-CAR-mediated activation at cell-cell contacts. CAR systems not only provide a powerful model to study early cell-cell interactions and the role of membrane topology and protrusions in signalling, but also offer insights that could guide the improved engineering of CAR T cells for more effective cancer targeting (Xiong et al., 2024). Our findings indicate that the HER2-CAR itself, LAT, Lck and CD45 do not localise preferentially to actin-rich protrusions prior to contact with a target cell. Within the first two minutes of target cell contact—preferentially initiated through actin-rich protrusions—CAR and LAT clustering, as well as ZAP-70 recruitment, are enhanced within these structures compared to other membrane regions. This is likely due to more efficient exclusion of CD45 from protrusions, as the kinase Lck shows no significant redistribution at the macroscale.

## Results

### Endogenously tagged signalling proteins are not enriched in actin-rich protrusions prior to T cell activation

To understand the contribution of protrusions to early signalling events in CAR-mediated T cell activation, we aimed to gain a better understanding of how key signalling proteins rearrange within T cell protrusions. As a first step, we analysed their nanoscale localisation under resting conditions. Previous studies have reported contradictory results on the localisation of signalling proteins in protrusive structures prior to activation in fixed T cells (Cai et al., 2022; Ghosh et al., 2020; Jung et al., 2021). Those reporting enrichment of TCR, Lck and LAT in protrusions suggested it may favour activation, as proteins confinement in microvilli may amplify early TCR signalling. We thus wanted to dissect the distribution of LAT, Lck and CD45 in actin-rich protrusions in an unperturbed living T cell using confocal and super-resolution STED microscopy, a purely optical imaging technique.

To avoid artefacts generated by protein overexpression, we knocked-in a Halo tag at the endogenous loci of ZAP-70, Lck, LAT or CD45 in Jurkat T cells (Fig. 1A). The successful tagging of the proteins was verified via western blot (Supplementary Fig. 1A), and the ability of the gene-edited knock-in (KI) cells to normally activate was assessed by measuring their IL-2 secretion (Supplementary Fig. 1B). While ZAP-70 displayed cytoplasmic localisation prior to activation, live-cell STED imaging of LAT, Lck and CD45 highlighted a strong plasma membrane localisation of all proteins that marked both the main body membrane and protrusions, without displaying any striking enrichment in protrusions (Fig.1A). Live-cell STED further highlighted local clustering of Halo-tagged Lck, LAT and CD45 at the plasma membrane, suggesting a certain degree of macromolecular assembly under resting conditions, consistent with previous reports (Lillemeier et al., 2010; Rossy et al., 2013; Sherman et al., 2011). To confirm actin enrichment and to later analyse the enrichment of signalling proteins in T cell protrusions, we developed an unbiased protrusions segmentation pipeline (Supplementary Fig. 2A). This approach allowed us to quantify protein *enrichment* by normalising the average intensity within protrusions to the average intensity across the full plasma membrane (Supplementary Fig. 2B). To account for possible sampling artefacts, such as increased signal in protrusions due to a higher membrane-to-pixel ratio, we compared the distribution of each protein with a SNAP-tagged CAAX motif (Moores et al., 1991), which serves as an evenly distributed plasma membrane marker. We calculated each protein’s *relative enrichment* by normalising its protrusion *enrichment* to that of the membrane marker in the same cell (Supplementary Fig. 2 B). A *relative enrichment* greater than 1 indicates enhanced localisation within protrusions, less than 1 indicates exclusion, and a value of 1 reflects no preferential localisation. T cell protrusions were highly enriched in actin, as shown by imaging the actin probe LifeAct-GFP (Fig. 1B) and quantifying its *enrichment* (Fig. 1C), which was significantly higher than the CAAX marker *enrichment*. On the other hand, analysis of confocal imaging of endogenously tagged Lck, LAT and CD45 yielded a *relative enrichment* around 1 (Fig. 1D-E) suggesting no preferential localisation of these proteins to actin-rich protrusions prior to activation. STED imaging further revealed the lack of protein enrichment in specific sub-domains within the protrusions in resting T cells (Fig. 1D).

**Figure 1:**
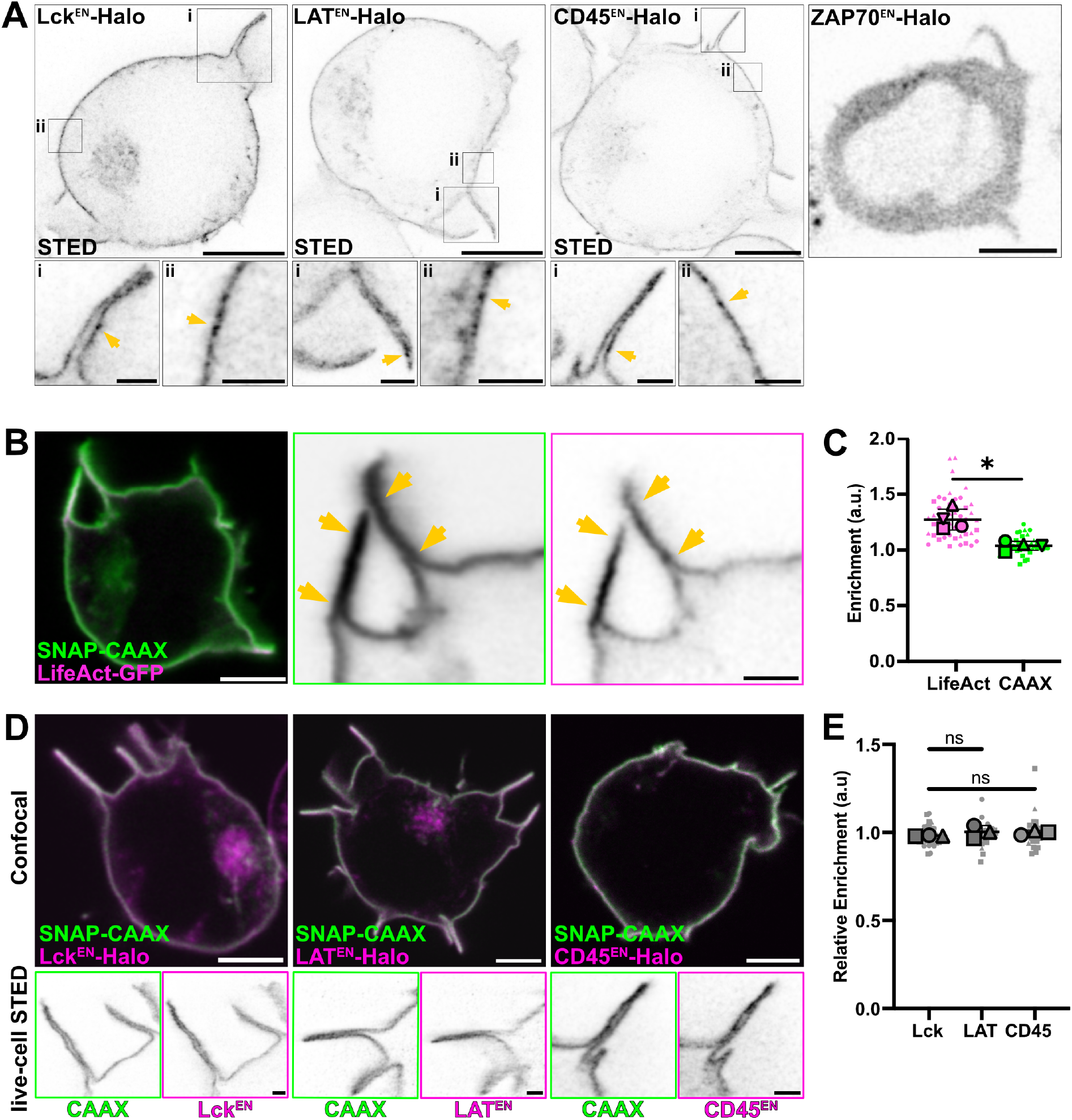
Live-cell STED imaging of endogenously tagged proteins revealed that signalling proteins are not enriched in actin-rich protrusions prior to T cell activation. **A)** Live-cell confocal and STED imaging of Jurkat T cells expressing endogenously Halo-tagged ZAP-70, Lck, LAT and CD45 labelled with CA-JFX_650_. Crops show protrusions (i) and main body membranes (ii). Arrows highlight the heterogeneous distribution of the proteins in the membrane. **B)** Live-cell imaging of Jurkat T cells expressing LifeAct-GFP and SNAP-CAAX (labelled with BG-JFX_650_). Arrows highlight protrusions enriched in F-actin. **C)** *Enrichment* of LifeAct-GFP or SNAP-CAAX in protrusions. In total, 45 cells from four independent experiments were analysed. Graph shows mean values, standard deviation (s.d.) error bars. P-value of paired t-test is 0.0142. **D)** Live-cell confocal (magenta and green) and STED images (inverted greyscale) of Jurkat T cells expressing Lck^EN^-Halo, LAT^EN^-Halo and CD45^EN^-LAP-Halo labelled with CA-JFX_650_ and SNAP-CAAX labelled with BG-JF_571_. **E)** *Relative enrichment* of LAT, Lck and CD45 in protrusions. In total, 32 cells (for Lck), 34 cells (for LAT) and 34 cells (for CD45) from three independent experiments were analysed. Replicates are shown as different shapes, and each small dot represents a single cell. Graph shows mean values, s.d. error bars. P-values of unpaired t-tests are 0.1411 (Lck/CD45) and 0.7628 (LAT/CD45). CA = chloroalkane (HaloTag substrate), BG = benzylguanine (SNAP-tag substrate). Scale bars, 5 μm (confocal overviews), 1 μm (crops and STED images).

### Contact with the target cell and clustering of CARs is enhanced by actin protrusions

As resting T cells showed no enrichment of the signalling proteins investigated in actin-rich protrusions, we next examined the localisation and dynamics of signalling proteins during early cell-cell interactions to assess the potential role of protrusions in early signalling. To trigger cell-cell interactions and T cell signalling, we employed a second-generation HER2-specific CAR containing an ectodomain derived from the HER2-specific FRP5 antibody and a CD28-CD3ζ endodomain (Ahmed et al., 2015). Live-cell STED imaging of HER2-CAR in resting Jurkat T cells highlights pre-clustering of the receptor (Supplementary Fig. 3A), however, quantification of the CAR *relative enrichment* highlighted no preferential localisation to actin-rich protrusions (Supplementary Fig. 3B). To follow the rearrangement of the receptor upon contact with a target cell, we co-cultured Jurkat T cells co-expressing GFP-tagged HER2-CAR and the membrane marker SNAP-CAAX with HER2^+^ SK-BR-3 cancer cells which exhibit elevated HER2 receptor levels on their cell surface (Dai et al., 2017) (Fig. 2A, Supplementary video 1).

**Figure 2:**
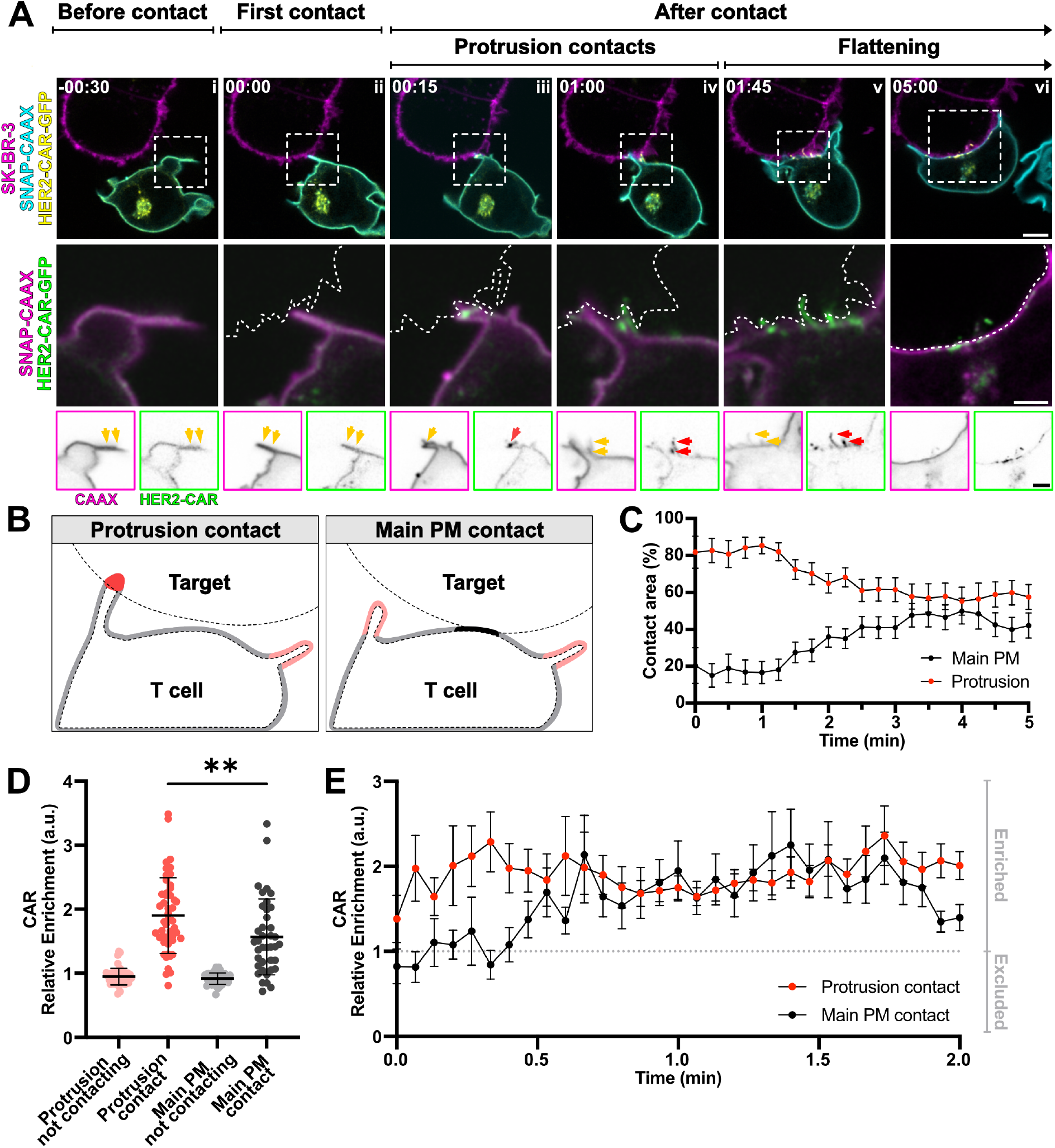
Contact with the target cell and early clustering of chimeric antigen receptors (CAR) is governed by protrusions. **A)** Time-lapse confocal imaging of a Jurkat T cell expressing HER2-CAR-GFP, SNAP-CAAX (labelled with BG-JF_571_) and interacting with a SK-BR-3 cell (labelled with CellMaskOrange). Dashed line describes the outline of the SK-BR-3 cell. Yellow arrows highlight protrusions, red arrows highlight the appearance of CAR clusters. **B)** Schematic representation of the four different plasma membrane regions segmented using the image analysis pipeline described in the Methods section and Supplementary Fig. 2. **C)** Percentage of contact area between the SK-BR-3 and the Jurkat T cell mediated by protrusions and by the main body membrane. n = 20 independent cell-cell interaction events captured with confocal microscopy at a 15s/frame rate are plotted. Graph shows mean values, standard error of the mean (s.e.m.) error bars. **D)** Mean *relative enrichment* of HER2-CAR-GFP in the four different regions described in B) during the first two minutes of the interaction with the SK-BR-3. n = 49 independent cell-cell interaction events interactions were analysed. P-value of paired t-test is 0.0094. Graph shows mean values, s.d. error bars. **E)** *Relative enrichment* of the CAR in the protrusions and main body membrane contacts over time. n = 29 captured with confocal microscopy at a 4s/frame rate are plotted. Graph shows mean values, s.e.m. error bars. BG = benzylguanine (SNAP-tag substrate). Scale bars, 5 μm (confocal overviews), 2 μm (crops).

Live-cell confocal microscopy allowed us to follow cell-cell interactions from the first contact until immunological synapse formation (Fig. 2A, Supplementary video 1). The first contact of the T cell with the target cell usually occured on actin-rich protrusions (Fig. 2Aii, Supplementary Fig. 4B). After the first contact, the HER2-CAR quickly clustered at protrusions (Fig. 2Aiii) and more protrusive structures polarised to the synapse (Fig. 2Aiv). These structures eventually collapsed (Fig. 2Av and vi), giving rise to a flattened immunological synapse. We could observe a similar behaviour in human primary CD4^+^ T cells (Supplementary Fig. 5, Supplementary video 2). Additionally, we often observed that projections from the SK-BR-3 cells contacted the Jurkat T cell and HER2-CARs were subsequentially accumulated at those contacts (Supplementary Fig. 4A and C, Supplementary video 3). This suggests that protrusions in general, and not exclusively from the T cell itself, can drive CAR accumulation at the interaction site. To evaluate the contribution of actin-rich protrusions and the main cell body to CAR clustering during early cell–cell interactions, we segmented these regions using our newly developed image analysis pipeline (Fig. 2B, Supplementary video 4). This allowed to quantify the involvement of protrusions and main cell body membranes in cell-cell contact over time (Fig. 2C). Our analysis revealed that actin-rich protrusions were the predominant feature at the T cell-target interface during the initial two minutes of interaction. To determine whether different plasma membrane regions—protrusions versus the main body—differ in their capacity to initiate clusters, we calculated the *relative enrichment* of the HER2-CAR in both regions, comparing contacting and non-contacting areas (Fig. 2B). During the first two minutes, all membranes in contact with the target showed enhanced CAR *relative enrichment* (Fig. 2D); however, *relative enrichment* was significantly higher in protrusion contacts in comparison to main body membrane contacts. In addition, quantitative analysis of the CAR *relative enrichment* after the first contact (set at t = 0s) revealed that CAR accumulated both rapidly and to a greater extent in protrusive contacts (Fig. 2E). A significant increase in CAR signal in protrusions was already noticeable 4s post-contact, while CAR enrichment in the main body membrane occurred only 30s post-contact. Overall, we demonstrate that protrusions promote the enrichment and clustering of CAR.

### CAR clusters in protrusions are activation hotspots

Taking advantage of our image analysis pipeline, we wanted to investigate whether CAR clusters in protrusions are activation hotspots by determining the dynamics of recruitment of downstream machinery to favour signal initiation. One of the earliest signalling events following CAR engagement and triggering is the recruitment of the tyrosine kinase ZAP-70 to the phosphorylated ITAMs of the cytoplasmic CD3ζ tail of the CAR. Therefore, ZAP-70 recruitment to the membrane is a reliable indicator of proximal CAR signalling (Gudipati et al., 2020). After its recruitment and activation by Lck phosphorylation, ZAP-70 phosphorylates the adaptor LAT (Zhang et al., 1998), that will scaffold a signalling hub for downstream signal propagation via calcium- and MAPK-dependent signalling pathways (Balagopalan et al., 2015). LAT clustering is thus another downstream readout of receptor activation.

To determine whether these CAR clusters are activation hotspots and whether increased CAR accumulation in protrusions correlates to an enhanced ZAP-70 recruitment, we expressed the HER2-CAR-GFP in a Jurkat T cell line expressing endogenously Halo-tagged ZAP-70 and followed the interaction with the target cell via confocal microscopy (Fig. 3A, Supplementary video 5). The recruitment of ZAP-70 to the CAR clusters immediately followed CAR clustering (Fig. 3Aii) after first contact (Fig. 3Ai). Although contacts via the main body membrane were uncommon, they also led to successful ZAP-70 recruitment. By employing our image analysis pipeline for automatic segmentation of the T cell membrane, we were able to calculate the *relative enrichment* of ZAP-70 at cell-cell contacts (Fig. 3B). ZAP-70 was more enriched at cell–cell contacts mediated by protrusions compared to those involving the main body membrane (Fig. 3B), suggesting that protrusions serve as more effective activation hotspots. We then asked whether enhanced ZAP-70 recruitment in protrusions was accompanied by faster and greater LAT clustering. For this, we employed a Jurkat T cell line expressing LAT^EN^-Halo and HER2-CAR-GFP and followed the first minutes of interaction with the target SK-BR-3 cells (Fig. 3C-D, Supplementary video 6). Shortly after target cell contact (Fig. 3Cii)— following CAR clustering and ZAP-70 recruitment—LAT rapidly clustered within actin-rich protrusions, typically between 8 and 12 seconds post-contact (Fig. 3D). In contrast, LAT clusters formation at main body interactions sites was delayed, with enrichment becoming apparent only at t = 32s post-contact (Fig. 3D). Overall, LAT clustering during the first minute of cell-cell interaction was significantly higher in protrusions than in the main body membrane (Fig. 3E), indicating that clustering of LAT is favoured in actin-rich protrusions. Interestingly, live-cell confocal microscopy highlighted clusters forming within cell-cell contacts in protrusions and then segregating from the HER2-CAR-positive cluster (Fig. 3Cii-iv). Live-cell STED imaging of cells expressing LAT^EN^-Halo and the membrane marker SNAP-CAAX confirmed the localisation of LAT clusters to the plasma membrane of protrusions and highlighted the segregation of the LAT clusters from the CAR clusters (Fig. 3F-G). Segregation of LAT from the membrane contact could be linked to the internalization and degradation of LAT (Balagopalan et al., 2007). Altogether, CAR clusters in protrusions are better activation hotspots than clusters derived from interactions via the main body as demonstrated by the accelerated kinetics of recruitment of downstream activation machinery.

**Figure 3:**
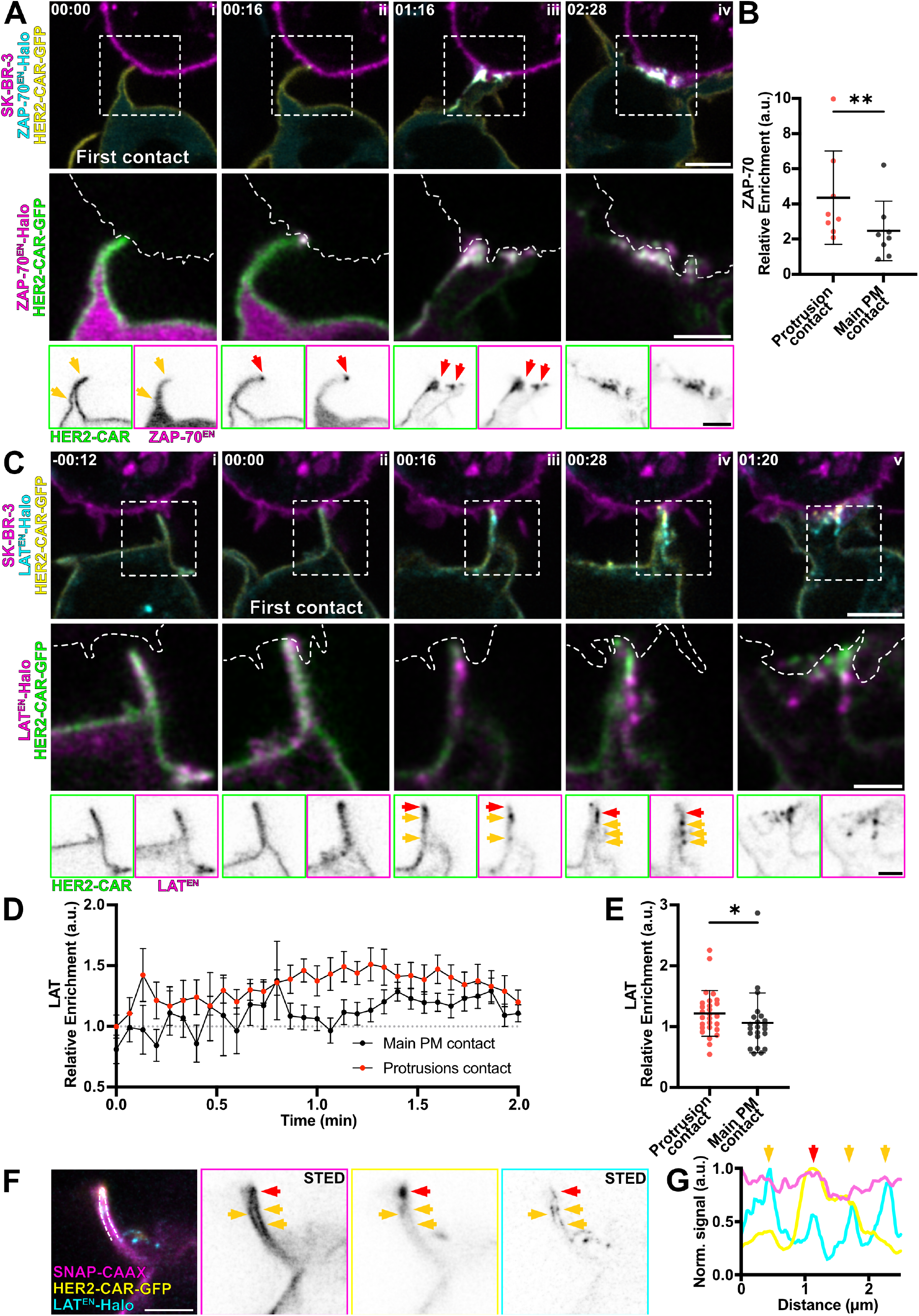
CAR clusters in protrusions are activation hotspots. **A)** Time-lapse confocal imaging of Jurkat T cells expressing HER2-CAR-GFP, ZAP-70^EN^-Halo labelled with CA-JFX_650_ interacting with a SK-BR-3 cell stained with CellMaskOrange. Dashed line describes the outline of the SK-BR-3 cell. Yellow arrows point at protrusions red arrows highlight ZAP-70 and CAR clusters, localised to protrusions. **B)** Mean *relative enrichment* of ZAP-70 in protrusions and main body membrane within the first minute of interaction with the target cell. n = 8 independent cell-cell interaction events were analysed. Graph shows mean values, s.d. error bars. P-value of paired t-test is 0.0020. *Relative enrichment* was computed as the *enrichment* in contacts divided by *enrichment* in protrusions or main body membranes not contacting the target. **C)** Time-lapse confocal imaging of Jurkat T cells expressing HER2-CAR-GFP, LAT^EN^-Halo labelled with CA-JFX_650_ interacting with a SK-BR-3 cell stained with CellMaskOrange. Dashed line describes the outline of the SK-BR-3 cell. Arrows highlight the formation of LAT^EN^ clusters in actin-rich protrusions and segregation from CAR clusters. Yellow arrows point at a LAT cluster colocalising with a HER2-CAR cluster and red arrows point at a LAT cluster segregated from a CAR cluster. **D)** *Relative enrichment* of LAT in actin-rich protrusions contacts and main body membrane contacts over time. n = 22 independent cell-cell interaction events are plotted. Graph shows mean values, s.e.m. error bars. **E)** Mean *relative enrichment* of LAT in protrusions and main body membrane within the first minute of interaction with the target cell. Graph shows mean values, s.d. error bars. P-value of paired t-test is 0.0367. **F)** Live-cell confocal and STED microscopy of HER2-CAR-GFP (confocal), SNAP-CAAX (STED) and LAT^EN^-Halo (STED) labelled with BG-JF_571_ and CA-JFX_650_ of the protrusion of a Jurkat T cell interacting with an unstained SK-BR-3. Yellow arrows point at a LAT cluster colocalising with a HER2-CAR cluster and red arrows point at a LAT cluster segregated from a CAR cluster. **G)** Line profile of the 3-colour image shown in F. Arrows correspond to the areas highlighted in D. CA = chloroalkane (HaloTag substrate), BG = benzylguanine (SNAP-tag substrate). Scale bars, 5 μm (confocal overviews), 2 μm (crops), 1 μm (STED).

### While Lck remains uniformly distributed, CD45 is excluded from target contacts

To better understand the mechanism behind the increased clustering of CAR and increased recruitment of signalling proteins in protrusions, we employed our quantitative live-cell imaging pipeline to assess the dynamics of localisation of negative (the phosphatase CD45) and positive (the tyrosine kinase Lck) regulators of CAR activation. According to the kinetic segregation model (Davis and van der Merwe, 2006; Xiao et al., 2022), the physical exclusion of the large phosphatase CD45 from close contacts would allow CAR to be phosphorylated by kinase Lck unopposed.

To monitor the dynamics of recruitment of the kinase Lck, we expressed the HER2-CAR in a gene-edited Jurkat T cell line expressing Lck^EN^-Halo (Fig. 4) and performed time-lapse confocal microscopy to assess any changes in the localisation of the protein upon contact with the target cell. Interestingly, we could not observe any macroscale-level rearrangement of Lck on the T cell membrane, whether in contact with the target cell or not (Fig. 4A, Supplementary video 7). While CAR clustering (Fig. 4Aiii) is prominent after first contact (Fig. 4Aii), LCK remains evenly distributed on the PM of T cells. Unbiased quantitative analysis highlights that Lck *enrichment* in various PM regions did not show any significant changes over time and upon cell-cell contact (Fig. 4B). No rearrangement of Lck^EN^-Halo in protrusions during the first minute of the interaction was detected (Fig. 4C). Furthermore, live-cell STED imaging of Lck^EN^-Halo cells expressing the membrane marker SNAP-CAAX could not highlight any nanoscale rearrangement of Lck within protrusions upon target cell contact (Fig. 4D). Taken together, this indicates that no macroscale-level rearrangement of Lck supported the heightened CAR activation at protrusions.

**Figure 4:**
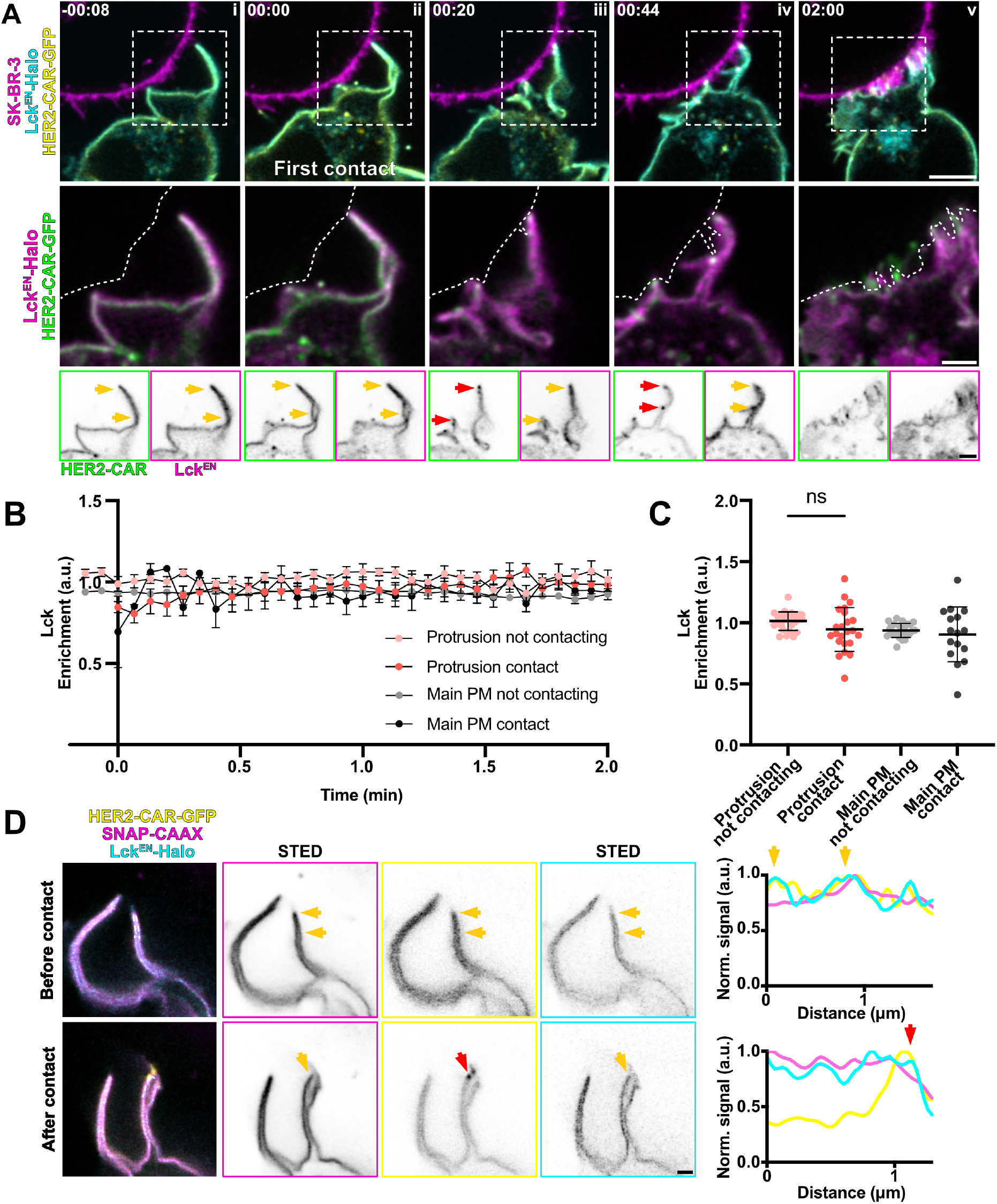
Lck remains evenly distributed on the plasma membrane of T cells upon CAR engagement. **A)** Time-lapse confocal imaging of Jurkat T cells expressing HER2-CAR-GFP, Lck^EN^-Halo labelled with CA-JFX_650_ interacting with a SK-BR-3 cell stained with CellMaskOrange. Dashed line describes the outline of the SK-BR-3 cell. Yellow arrows highlight protrusions and the lack of rearrangement of Lck at the macroscale, while red arrows point at CAR clusters at protrusions upon contact with the target cell. **B**) *Enrichment* of Lck in the four different membrane regions segmented as described in Fig. 2B. n = 26 independent cell-cell interaction events are plotted. Graph shows mean values, s.e.m. error bars. **C)** Mean *enrichment* of Lck^EN^ in protrusion and main body membrane contacting or not contacting the target during the first minute of the interaction. n = 23 independent cell-cell interaction events were analysed. Graph shows mean values, s.d. error bars. P-value of paired t-test is 0.0939 for protrusion regions and 0.4764 for body membrane regions. **D)** Live-cell confocal and STED microscopy of HER2-CAR-GFP (confocal), SNAP-CAAX (STED) and Lck^EN^-Halo (STED) labelled with BG-JF_571_ and CA-JFX_650_ of a protrusion of a Jurkat T cell before and after interacting with an unstained SK-BR-3. The section corresponding to the line profile is shown in the three-colour image. Yellow arrows highlight protrusions and the lack of rearrangement of Lck at the macroscale, while the red arrow points at the CAR cluster in a protrusion upon contact with the target cell. CA = chloroalkane (HaloTag substrate), BG = benzylguanine (SNAP-tag substrate). Scale bars, 5 μm (confocal overviews), 2 μm (crops), 1 μm (STED).

We then performed the same experiment with gene-edited Jurkat T cells expressing CD45^EN^-Halo (Fig. 5). Strikingly, as the T cell makes first contact with the target cell (Fig. 5Aii), CD45 became excluded from the CAR cluster formed in protrusions (Fig. 5Aiii, Supplementary video 8). CD45 signal intensities markedly decreased at the sites of initial protrusion contact and CAR clustering (Fig. 5Aii, iii and iv, line profiles). When analysing the *relative enrichment* of CD45 over time (Fig. 5B), we could observe that the exclusion of the protein in contacts with the target cell was immediate (t = 0 s). On the contrary, decrease of CD45 signal at main body membrane contacts, only became apparent at later stages of cell-cell interaction (t = 40 s). Accordingly, the average *relative enrichment* of CD45 at protrusion contacts was significantly lower than that at main body membrane contacts (Fig. 5C). While CD45 exclusion at protrusion-mediated contacts was only modestly apparent by confocal microscopy, live-cell super-resolution STED imaging provided a more detailed view—resolving individual protrusions and membrane contours—clearly highlighting CD45 exclusion at sites of CAR clustering (Fig. 5D–E). Altogether, this data indicates that CD45 exclusion might be faster and more efficient at contacts created through protrusive structures, explaining the increased clustering and activation of CARs in protrusions.

**Figure 5:**
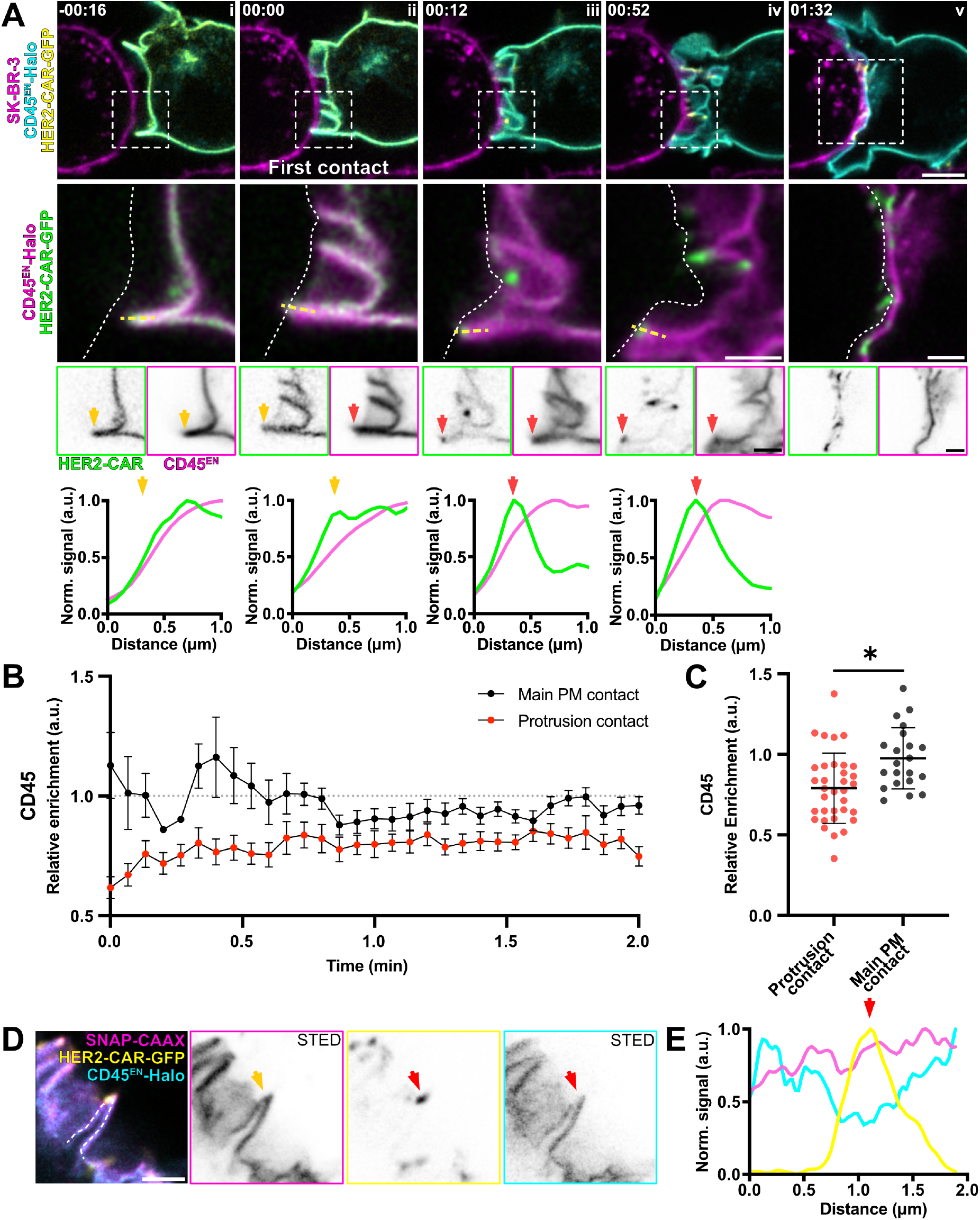
CD45 is excluded from contacts with the target cell and the exclusion is more efficient in actin-rich protrusions. **A)** Time-lapse confocal imaging of Jurkat T cells expressing HER2-CAR-GFP and CD45^EN^-Halo (labelled with CA-JFX_650_) interacting with a SK-BR-3 cell stained with CellMaskOrange. Dashed line describes the outline of the SK-BR-3 cell. Yellow arrows highlight a protrusion displaying non-preferential localisation of CD45 or the CAR prior to contact with the target cell and red arrows highlight a membrane domain with decreased CD45 signal and CAR enrichment. Box profiles show HER2-CAR and CD45 signal along the dashed yellow line. **B)** *Relative enrichment* of CD45 in actin-rich protrusions contacts and main body membrane contacts over time. n = 34 independent cell-cell interaction events are plotted. Graph shows mean values, s.e.m. error bars. **C)** Mean *relative enrichment* of CD45 in protrusions and main body membrane within the first minute of interaction with the target cell. Graph shows mean values, s.d. error bars. P-value of paired t-test is 0.0169. **D)** Live-cell confocal and STED microscopy of HER2-CAR-GFP (confocal), SNAP-CAAX(STED) and CD45^EN^-Halo labelled with BG-JF_571_ and CA-JFX_650_ in a Jurkat T cell interacting with an unstained SK-BR-3. Red arrow points at a HER2-CAR cluster formed at a cell-cell contact to highlight the dramatic decrease of CD45 signal intensity. **E)** Line profile of the 3-colour image shown in F. Arrow corresponds to the area highlighted in D. CA = chloroalkane (HaloTag substrate), BG = benzylguanine (SNAP-tag substrate). Scale bars, 5 μm (confocal overviews), 2 μm (crops), 1 μm (STED).

Next, we wanted to test whether exclusion of CD45 depends on CAR signalling or is simply driven by the physical interaction between CAR and antigen. The first indication that this may be solely a physical process came from the observation that CAR *relative enrichment* increased around t = 4 s (Fig. 2Aiii, C), while CD45 exclusion was evident upon first contact at t = 0 s (Fig. 5Aii-B). To assess whether the process is independent of CAR signalling, we expressed a truncated version of the CAR in the CD45^EN^-Halo Jurkat T cell line (Fig. 6). The truncated CAR only harbours the extracellular HER2-binding domain, a transmembrane domain and a GFP tag and is thus unable to signal. We assessed the rearrangement of the truncated CAR and CD45 in time-lapse experiments upon contact with the target cell (Fig. 6A, Supplementary video 9). We found that CD45 exclusion at contact sites happened instantaneously (Fig. 6Aii, t = 0 s), followed by a clear clustering of the truncated CAR (Fig. 6Aiii, t = 16 s). This observation mirrors that seen with CARs with an intact signalling domain. However, despite the formation of initial contacts, the membrane of most cells expressing the truncated CAR did not flatten to form a mature synapse (Fig. 6Av), indicating incomplete activation. Further, comparison of the *relative enrichment* of CD45 at protrusion contacts stabilised by the intact and truncated CAR, we could not observe any significant difference in CD45 exclusion (Fig. 6B). All together, these results indicate that CD45 exclusion at cell-cell contacts is CAR signalling-independent and precedes CAR clustering and recruitment of downstream machinery.

**Figure 6:**
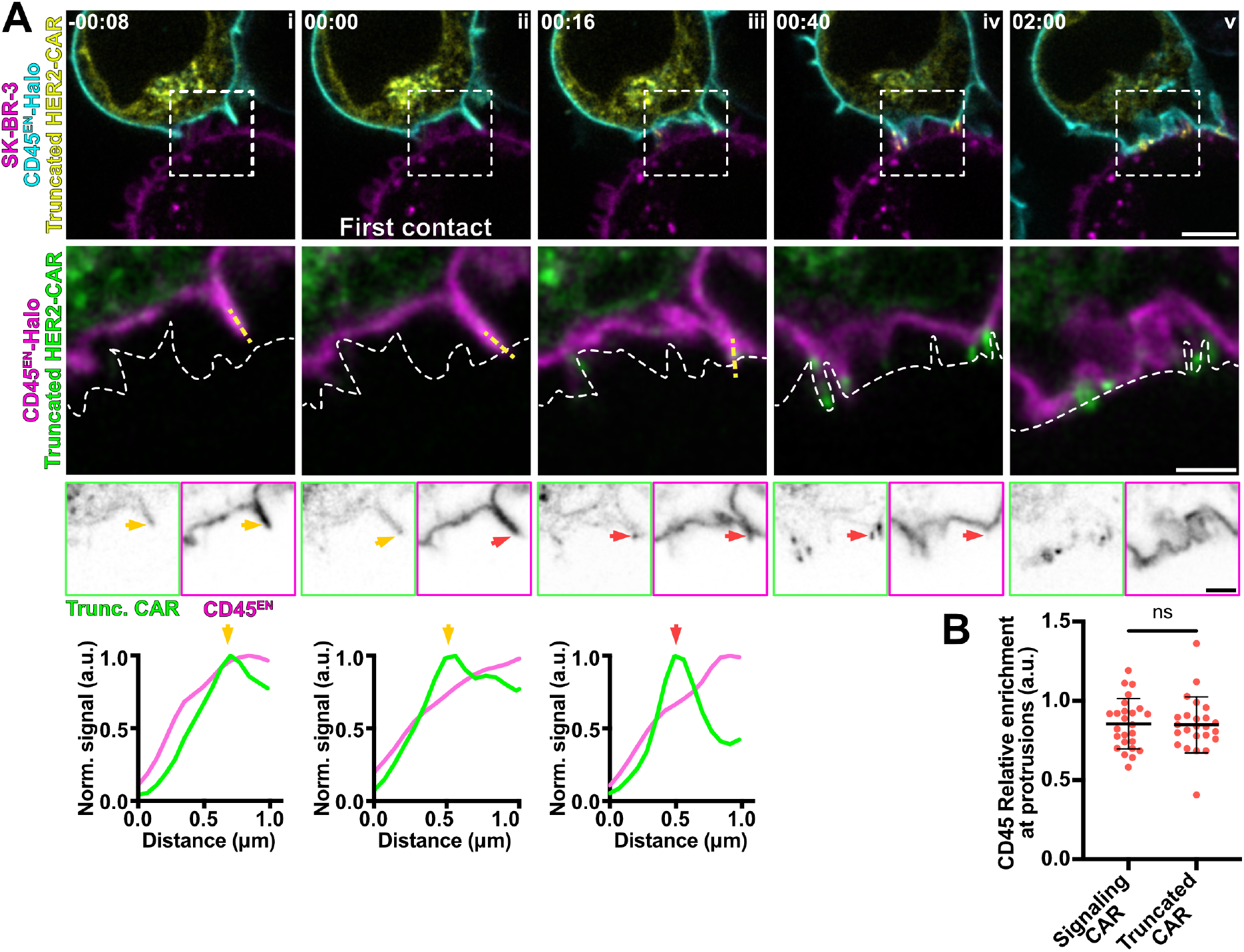
CD45 is excluded from protrusion contacts with the target cell independently of signalling. **A)** Time-lapse confocal imaging of Jurkat T cells expressing a truncated HER2-CAR-GFP and CD45^EN^-Halo labelled with CA-JFX_650_ interacting with a SK-BR-3 cell stained with CellMaskOrange. Dashed line describes the outline of the SK-BR-3 cell. The truncated HER2-CAR (Trunc. CAR) lacks form the co-stimulatory and signalling domains. Yellow arrows highlight a protrusion displaying non-preferential localisation of CD45 or the truncated CAR prior to contact with the target cell and red arrows highlight a membrane domain with decreased CD45 signal and truncated CAR enrichment. Line profiles show truncated HER2-CAR and CD45 signal along the dashed yellow line. **B)** Mean *relative enrichment* of CD45 in protrusion contacts during the first minute of interaction with the target cell mediated by the signalling CAR or by the truncated HER2-CAR. (Signalling HER2-CAR experiment) n = 18 independent cell-cell interaction events (truncated HER2-CAR) n = 24 independent cell-cell interaction events. P-value of unpaired t-test is 0.8954. CA = chloroalkane (HaloTag substrate. Scale bars, 5 μm (confocal overviews), 2 μm (crops).

## Discussion

In this study, we establish a quantitative, dynamic framework that captures macromolecular rearrangements in relation to membrane topography on the surface of T cells—from first cell-cell contact to immunological synapse formation—all within living, unperturbed cells. This approach offers unprecedented insight into the spatial and temporal coordination of signalling events in their native context. Our findings indicate that prior to activation, key signalling machinery lacks any enrichment in actin-rich protrusions (Fig. 7B). Here, we define actin-rich protrusions as plasma membrane extensions enriched in F-actin (Fig. 1B-C), which enabled us to assess the general function of this kind of structures. However, upon contact with the target cell, we observed a marked increase in CAR and LAT clustering, enhanced ZAP-70 recruitment and CD45 exclusion within protrusions compared to the main cell body membrane regions (Fig. 7D vs Fig. 7C). Overall, this study reveals that actin-rich protrusions are privileged sites for the initiation of CAR-mediated T cell activation.

**Figure 7:**
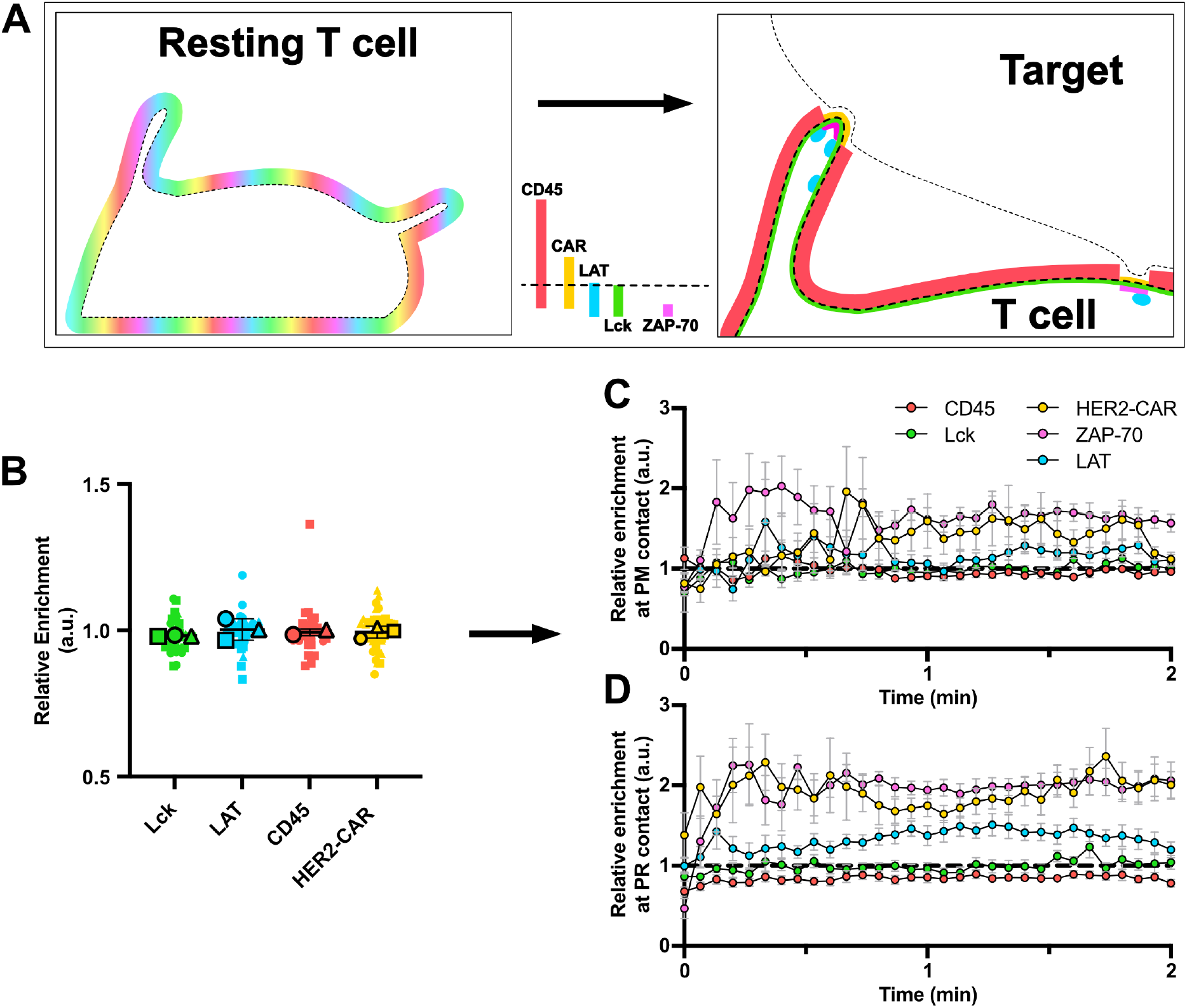
A quantitative framework for the dynamic rearrangement of signalling proteins on the T cell surface. **A)** Schematic representation of the localisation of the HER2-CAR, Lck, ZAP-70, LAT and CD45 before and at early contact with the target cell. CD45 is excluded from plasma membrane regions at close cell–cell contacts formed both at protrusions and the main body of the T cell, with exclusion being more pronounced at protrusions. This promotes enhanced CAR clustering and activation in protrusions, shown by earlier and greater signal increase of CAR, ZAP-70, and LAT within protrusions at contact sites. **B)** Quantification of the *relative enrichment* of Lck, LAT, CD45 and the HER2-CAR in resting conditions. Results around 1 indicate no localisation preference to protrusions in comparison to the membrane marker. **C and D)** *Relative enrichment* at C) Main body membrane and D) protrusion membrane contacts with the target cell. t = 0 min indicates the first contact with the target cell. Graphs show mean values, s.e.m. error bars.

In resting T cells, we observed that HER2-CAR, Lck, LAT and CD45 do not exhibit preferential localisation to protrusions (Fig. 1E, Supplementary Fig. 3B, Fig. 7B). This differs from a previous report suggesting that Lck and LAT may be pre-assembled in protrusive structures known as “microvilli” (Ghosh et al., 2020). Similar discrepancies have been noted in studies of TCR localisation, likely due to differences in imaging methodologies. For example, while some studies reported TCR enrichment in protrusions (Ghosh et al., 2020; Jung et al., 2016) others found no such enrichment (Cai et al., 2022). To address these discrepancies, we sought to characterize the resting distribution of signalling proteins in living cells using a purely optical super-resolution microscopy technique such as STED (Lukinavičius et al., 2024). By doing so, we aimed to distinguish between pre-existing spatial arrangements and dynamic protein redistributions that occur specifically upon target engagement. Our findings strongly support the latter and highlight the importance of live-cell imaging for accurately capturing the native organisation of signalling components.

Upon contact with the target cell, actin-rich protrusions became the primary structure mediating early interaction at the interface (Fig. 2C). Even after a partial flattening of the T cell, we could observe some protrusive interdigitations with the breast cancer cell (Fig. 2A iv), which have also been previously reported (Sanderson and Glauert, 1979). It is worth noting that cell-cell interactions governed by protrusions might be more pronounced in the CAR-mediated system, as CAR engagement leads to hyper-stabilization of protrusive structures (Beppler et al., 2023). By imaging the HER2-CAR rearrangement in a Jurkat T cell interacting with the target, we could monitor how the different membrane regions contribute to close contact formation and receptor recruitment. The protrusive behaviour of Jurkat T cells is identical to the one of peripheral blood-derived CD4+ primary T cells, suggesting analogous mechanisms (Supplementary Fig. 5). We found that, although CARs accumulated at both protrusions and main body membrane interacting with the target, CAR *enrichment* was consistently higher in protrusions (Fig. 2D). This suggests that protrusions are more efficient at initiating close contacts. In some cases, we could observe how target cell protrusions engaged the T cell main body membrane and triggered CAR clustering (Supplementary Fig. 4A). This observation aligns with previous studies showing that dendritic F-actin structures facilitate T cell signalling and priming (Leithner et al., 2021). Since T cell protrusions exert pushing forces that promote close apposition of membranes (Sage et al., 2012; Sanderson and Glauert, 1979), it is plausible that protrusions from the target cell may be able to do the same.

The increased HER2-CAR clustering in actin-rich protrusions was accompanied by an enhanced ZAP-70 accumulation at protrusions (Fig. 3B), indicating that these clusters functioned as activation hotspots from the moment of their formation. This was followed by faster (Fig. 3D) and more pronounced (Fig. 3E) LAT clustering at the same sites. Although this result could be simply explained by the increased HER2-CAR/ZAP-70 signal at protrusions, the architecture of the protrusion itself might be relevant for enhancing the immune response, as we speculate that negative curvature may favour accumulation of signalling proteins by reducing diffusion within membranes. We were able to show that the primary mechanism by which actin-rich protrusion promote increased CAR accumulation and activation is their ability to exclude CD45 from the interaction site faster and in a higher magnitude than the main body membrane (Fig. 5). CD45 has a large and rigid extracellular domain (Chang et al., 2016a) that extends beyond the distance spanned by the intercellular HER2-CAR/HER2 complex. According to the kinetic segregation model, receptor triggering would occur because of size-dependent exclusion of CD45 from close contacts in which the receptor would encounter its ligand (Davis and van der Merwe, 2006). A recent study has reported that CAR activation is dependent on the length of the CAR/ligand complex (Xiao et al., 2022), suggesting that kinetic segregation of CD45 is also a mechanism for CAR activation. Here we observe that protrusive structures have a greater ability to exclude CD45 than main body membranes (Fig. 5C) and the exclusion in protrusions occurs right upon contact, while exclusion at main body membranes occurs later in time (Fig. 5B). This early exclusion is CAR signalling-independent (Fig. 6) and precedes CAR recruitment to the contact (Fig. 2D vs Fig. 5C). The enhanced segregation in protrusions was enough to provide a nanoscale environment that allowed CAR activation even in the absence of macroscale rearrangement of the positive regulator of activation, Lck (Fig. 4). Lack of Lck rearrangement in CAR-mediated interaction is possibly explained by the absence of CD4-bound Lck brought in close proximity of the CAR by MHC-CD4 interactions (Wu et al., 2023; Zhang et al., 2025).

Combining live-cell super-resolution imaging of endogenously tagged proteins with quantitative analysis of membrane topography and protein dynamics allowed us to understand the role of membrane topography in cell-cell interactions. This study focuses on understanding how actin-rich protrusion drive early cell-cell interactions and signalling in CAR-mediated activation. The established cell lines and methodologies could be employed to investigate fundamental questions about the role of membrane topography in TCR-mediated activation and its function at later timepoints of interaction with a target.

## Methods

### Mammalian cell culture

Jurkat T cells (Cat # ACC 282 DSMZ) and SK-BR-3 cells (Cat # ACC 736 DSMZ) were grown in a humidified incubator at 37°C with 5% CO_2_ in Roswell Park Memorial Institute medium (RPMI, Gibco) supplemented with 10% fetal bovine serum (Corning), 100 U/L penicillin and 0.1 g/L streptomycin (FisherScientific). Normal human peripheral blood CD4+ primary T cells (Cat # SER-CD4-PLUS-T-F-ZB, BioCat) were grown in ImmunoCult-XF T cell expansion medium (Stemcell Technologies) supplemented with 10% fetal bovine serum, 100 U/L penicillin, 0.1 g/L streptomycin and 10 ng/mL human Recombinant IL-2 (Stemcell Technologies) at 37ºC with 5% CO_2_.

For transient transfection of plasmids encoding for SNAP-CAAX, HER2-CAR-GFP, HER2-CAR-Halo or truncated HER2-CAR-GFP into Jurkat T cells, a NEPA21 electroporation system was used (Nepa Gene). 5 million cells were washed twice with Opti-MEM (Gibco), resuspended in 90 µl Opti-MEM and mixed with 10 µg of DNA in an electroporation cuvette with a 2-mm gap (Fisher Scientific). The electroporation reaction consists of two poring pulses (150 V, 5 ms length, 50 ms interval, with decay rate of 10% and + polarity) and five consecutive transfer pulses (20 V, 50 ms length, 50 ms interval, with a decay rate of 40% and ± polarity). For transient transfection of SNAP-CAAX and HER2-CAR-GFP in primary T cells, CD4+ T cells were activated with ImmunoCult Human CD3/CD28 T cell Activator (Stemcell Technologies) following manufacturer’s instructions. After 48h, 3 million cells were electroporated using the NEPA21 system. The electroporation reaction consists of two poring pulses (175 V, 5 ms length, 50 ms interval, with decay rate of 10% and + polarity) and five consecutive transfer pulses (20 V, 50 ms length, 50 ms interval, with a decay rate of 40% and ± polarity).

### Generation of plasmids for overexpression and gene editing

For all plasmids harboring a Halo-encoding sequence, the HaloTag sequence was taken from pH6HTC His6HaloTag® T7 (Promega, JN874647). For all plasmids harboring a SNAP-encoding sequence, the SNAP-tag sequence was taken from the pSNAPf (Addgene plasmid, #58186). For all plasmids harboring a GFP-encoding sequence, the GFP sequence was taken from pEGFP-N1 (Addgene plasmid, #6085-1). All plasmids were verified through sequencing.

### Cloning of overexpression plasmids

To generate the SNAP-CAAX encoding plasmid, the GFP tag was replaced with a SNAP tag in the GFP-CAAX plasmid (Addgene plasmid, #86056). GFP-CAAX was digested with NheI and EcoRI and the SNAP sequence amplified from pSNAPf plasmid and ligated into the digested plasmid as an NheI-EcoRI fragment. Sequences of all primers used for fragment amplification are provided in Supplementary Table 1.

For the generation of the HER2-CAR-GFP, the truncated HER2-CAR-GFP and the LifeAct-GFP plasmids a similar strategy was used. A double strand DNA fragment with the desired sequence was synthesized (IDT) and cloned into the EcoRI/BamHI linearised eGFP-N1 vector (Clontech) via Gibson’s assembly. A detailed list of all gBlocks used for cloning is provided in Supplementary Table 2.

For the generation of the HER2-CAR-Halo plasmid, the HER2-CAR-GFP plasmid was digested with BamHI/NotI. A BamHI/NotI Halo fragment was then ligated into the linearised vector.

### Cloning of guide RNA and homology repair (HR) plasmids

Designing of plasmids and cloning was carried out as explained in detail in (Adarska et al., 2025). All guide RNAs were designed using Benchling (https://www.benchling.com). Single strand sense and antisense oligoes containing the guide sequence and harbouring BpII sites were synthesized and cloned via oligo annealing into the SpCas9 pX330 plasmid (addgene plasmid #42230) (Cong et al., 2013), which was previously linearised with BpII (Life Technologies). A detailed list of guide RNA sequences is provided in Supplementary Table 3. To prevent re-cutting from the Cas9, the protospacer-adjacent motif (PAM) site in the HR plasmids was mutated.

#### HR plasmid CD45/ZAP-70^EN^-LAP-Halo-polyA-G418^r^

The HR plasmid consists of ∼600 bp homology arms in a High Copy Amp plasmid (pMB1) backbone (Twist Bioscience). A glycine-serine linker and BamHI/EcoRI sites were added between the two homology arms for cloning of tags and a resistance cassette for antibiotic selection of positively edited cells. The coding sequences of a localisation and affinity purification (LAP) linker (Cheeseman and Desai, 2005), the HaloTag and a small 2xALFA epitope-tag were obtained from the ARF3-LAP-Halo-2xALFA-LoxP-G418^r^-LoxP HR plasmid (Wong-Dilworth et al., 2023) as a NheI/EcoRI fragment. The polyA sequence and the G418 resistance cassette were obtained via PCR amplification from theARF1-Halo homology repair plasmid (Bottanelli et al., 2017).

#### HR plasmid Lck ^EN^-Halo-polyA-G418^r^

The HR plasmid consists of ∼1000 bp homology arms in a High Copy Amp plasmid (pMB1) backbone (Twist Bioscience). BamHI and EcoRI sites were added for insertion of the Halo tag sequence and the resistance cassette. A short glycine-serine linker was added between two BamHI sites. The HaloTag sequence was obtained from the AP1µA^EN^-Halo-ALFA-polyAG418 plasmid (Stockhammer et al., 2024) as a BamHI/EcoRI fragment.

#### HR plasmid LAT ^EN^-Halo-polyA-G418^r^

The HR plasmid consists of ∼1000 bp homology arms in a High Copy Amp plasmid (pMB1) backbone from Twist Bioscience. A glycine-serine linker and a NheI and EcoRI sites were added between the two homology arms for insertion of the Halo tag sequence and resistance cassette. The HaloTag, a small 2xHA epitope tag, a polyA and the G418 resistance cassette sequences were amplified from the 1ARF1-Halo HR plasmid (Bottanelli et al., 2017).

### Generation of CRISPR Knock-In cell lines

For generation of Jurkat Knock-In cell lines, 4 million cells were washed twice with Opti-MEM (Gibco), resuspended in 90 µl Opti-MEM and mixed with 5 µg of HR plasmid and 5µg of sgRNA/Cas9 plasmid in an electroporation cuvette with a 2-mm gap. Electroporation was performed as described in “Mammalian cell culture”. G418 was added to the cells 3 days after transfection at a concentration of 3 mg/mL (G418) and media was exchanged every 2-3 days until selection was complete (approximately after 7 days). 0.5% brightest cells for the endogenously tagged Lck, LAT or CD45 cell lines and 0.01% brightest for the ZAP-70 cell line were isolated via Fluorescence-Activated Cell Sorting (FACS).

### SDS page and western blot

2 million cells were lysed in 200 µl of RIPA buffer [150nM NaCl (Roth), 0.1% Triton X-100 (Sigma-Aldrich), 0.5% sodium deoxycholate (Sigma-Aldrich), 0.1% SDS (CarlRoth) and 50mM Tris-Hcl (CarlRoth) pH 8.0, 1x protease inhibitor cocktail (Roche)]. Cells lysates were clarified by centrifugation for 10 min at 4ºC and 14,000 g and mixed 1:1 with 2x Laemmli buffer [4% SDS, 20% Glycerol (CarlRoth), 120nM TRIS-Hcl pH 6.8, 0.02%, Bromophenol (VWR Chemicals) and 5% β-mercaptoethanol (CarlRoth)]. Cell lysates were then loaded on 4-12% SDS-polyacrylamide gels (Life Technologies). After electrophoresis, proteins were transferred to a nitrocellulose membrane (Amersham, Fisher Scientific GmbH) via wet blotting. Membranes were blocked with 5% milk powder (Roth) and 1% BSA (Roth) in PBST [1xPBS (phosphate buffer saline, Corning), 0.5% Tween 20 (Roth)] and incubated with primary antibodies at 4ºC overnight. For detection, secondary horseradish peroxidase-coupled antibodies were used. For protein detection, Supersignal West Pico Plus chemiluminescent substrate (Thermo Scientific) was used. When necessary, membranes were stripped with stripping buffer containing 0.2M glycine pH 2.3, 35mM SDS and 0.1% Tween 20 (Roth). The antibodies employed are listed in Supplementary Table 4.

### IL-2 ELISA

200.000 cells were seeded in each well of an 8-well chambered cover glass (No. 1.5, Cellvis) coated with anti-human CD28 antibody (Biolegend) and Anti-Human CD3 antibody (Invitrogen) and incubated overnight at 37ºC and 5% CO_2_. Supernatants were harvested and IL-2 secretion was assessed using the ELISA MAX Deluxe Set Human IL-2 kit (Biolegend) following manufacturer’s indications. Absorbance of 560 and 405 nm light was measured on the Spark multimode microplate reader (TECAN).

### Labelling for live-cell imaging

For live-cell imaging, cells were labelled with Halo and/or SNAP substrates indicated in the figure legends at a concentration of 1 µM for 1 h at 37ºC in culture media. Cells were washed in growth media at 37 °C for at least 1 h to allow efflux of unbound dye. For imaging cells in resting conditions, 100.000 cells were seeded on 8-well chambered coverglass dishes (No. 1.5, Cellvis).

For imaging of SK-BR-3/Jurkat T cell interactions, 25.000 SK-BR-3 cells were seeded on 8-well chambered coverglass dishes and cultured overnight. SK-BR-3 cells were stained before imaging with the CellMask Orange plasma membrane stain (Invitrogen) for 5 min at 37ºC.100.000 Jurkat T cells expressing the HER2-CAR-GFP and the protein of interest, were added to the wells and imaged immediately after. Live-cell imaging was performed in live-cell imaging solution [FluoBrite DMEM (Gibco) supplemented with 10% FBS, 20 mM HEPES (Gibco) and 1x GlutaMAX (Gibco)].

### Imaging and image processing

Microscopy data was collected on an expert line Abberior STED microscope using the Imspector software from Abberior instruments (Version 16.3). The microscope is equipped with 485 nm, 561 nm and 640 nm excitation lasers. For two-colour STED experiments both JF_571_ (Grimm et al., 2020) and JFX_650_ (Grimm et al., 2021) dyes were depleted with a 775 nm depletion laser. The detection windows were set to 571 to 630 nm and 650 to 756 nm. Multi-colour STED images were recorded sequentially line by line. For three-colour confocal imaging, detection windows were set to 498 to 540, 590 to 625 and 660 to 757 nm. If required for quantitative analysis, laser power was kept constant between images. For live-cell confocal imaging the pixel size was set to 70 nm and for STED imaging to 30 nm. Live-cell imaging was performed at 37ºC.

To reduce noise, confocal images were background subtracted (rolling ball algorithm, radius of 50 pixels) and gaussian blurred (σ = 1 pixel) using Fiji (ImageJ 2.7.0, (Schindelin et al., 2012)). STED images in Fig. 1D, 3F, 4D and 5E were deconvolved using Richardson–Lucy deconvolution implemented in the python microscopy PYME package (https://python-microscopy.org). Line profiles shown in Fig. 3E, 4D and 5F were obtained using Fiji by drawing a line (with 3-pixel width) alongside the protrusion membrane on gaussian blurred (σ = 1 pixel) STED images. Line profiles on Fig. 5A and 6A were obtained by drawing a line (with 3-pixel width) alongside the protrusion on background subtracted (rolling ball algorithm, radius of 50 pixels) and gaussian blurred (σ = 1 pixel) confocal images. The line profile data was then normalised and plotted using GraphPad Prism (GraphPad Software, https://www.graphpad.com).

### Image Quantification

#### Segmentation of protrusions and main body membrane

Segmentation and quantification of Jurkat T cell membrane structures were performed using a custom image analysis pipeline (Supplementary Fig. 2A) implemented in CellProfiler (Stirling et al., 2021). 2-channel pictures of the Jurkat T cell membrane were decomposed into individual .tiff files using Fiji and loaded into CellProfiler. The two channels were then averaged and blurred with a gaussian filter (σ = 2 pixels). Segmentation on data from Fig. 3B was performed utilising only the HER2-CAR-GFP channel. Segmentation continued in two parallel streams: identification of the inner body/neighborhood and identification of membrane structures. For inner body detection, dark holes from the membrane image were enhanced using the rolling ball algorithm in the inverted image. These were thresholded and filtered by area, and internal holes were filled. The resulting inner body image was dilated by 9 pixels to cover the main body membrane, and by 35 pixels to define the cell neighborhood. For membrane detection, neurite-like structures were enhanced using the “tubeness” enhancement method and then thresholded. To obtain initial protrusion candidates, the 9-pixel-dilated inner body mask was subtracted from the thresholded neurite structures; negative values were set to 0. The final protrusion mask was obtained by filtering out structures outside of the neighborhood size filtering for discarding single-pixel objects. The main body membrane mask was obtained by subtracting the inner body and the protrusion masks from the thresholded neurites. The whole membrane mask was obtained by merging the protrusion and the main body membrane masks.

#### Segmentation of membrane contacts

To distinguish between target-contacting and not contacting regions of the Jurkat T cell membrane, SK-BR-3 cell masks was first generated with Fiji. The SK-BR-3 channel was background subtracted (rolling ball radius = 50 pixels), gaussian blurred (σ = 2 pixels). and thresholded with the “Moments” algorithm applied to the last frame of the stack. The SK-BR-3 mask, along with the HER2-CAR and protein of interest channels were loaded into CellProfiler. Protrusions and main body membranes were segmented as described above.Contacting regions were determined by applying an “and” operation between SK-BR-3 and the Jurkat T cell protrusions or main body membrane masks. Non-contacting regions were obtained by subtracting the contacting masks from their respective full masks.

To estimate the contribution of both membranes to the contact with the target in Fig. 2C, Jurkat T cells expressing HER2-CAR-GFP and SNAP-CAAX were imaged interacting with SK-BR-3 cells at a framerate of 15/seconds per frame. The areas occupied—in pixels—by the protrusion and main body membrane contacts were quantified and expressed as percentages of total contact area.

#### Enrichment and relative enrichment

Signal intensity values for each protein were normalised to their maximum values across the whole plasma membrane mask. The parameter “*enrichment*” (Supplementary Fig. 2B) of a protein in a membrane region was defined as

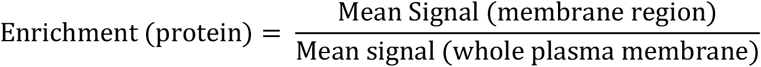

For Fig. 1D, dual-color confocal images of Lck^EN^-Halo/SNAP-CAAX, LAT^EN^-Halo/SNAP-CAAX and CD45^EN^-Halo/SNAP-CAAX were used, and the *relative enrichment* (Supplementary Fig. 2B) was computed as

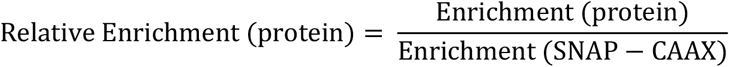

For Fig. 2E, three-color confocal time lapses of a HER2-CAR-GFP/SNAP-CAAX expressing Jurkat T cell interacting with a SK-BR-3 cell were obtained with a frame rate of 4s/frame. Time zero was set as the first frame in which a contact was detected by the pipeline. The CAR *relative enrichment* was computed as above. Fig. 2D reports the average of the first two minutes using data collected at both 4 s and 15 s intervals.

For Figs. 3B, 3D, 4B, 4C, 5B, 5C, and 6B, Jurkat cells expressing HER2-CAR-GFP or the truncated HER2-CAR and the endogenously tagged protein of interest were imaged during interactions with SK-BR-3 cells at 4 s/frame. For Fig. 3D and 5B, the *relative enrichment* of the protein in a membrane contact region (either protrusion or main body membrane) was computed as

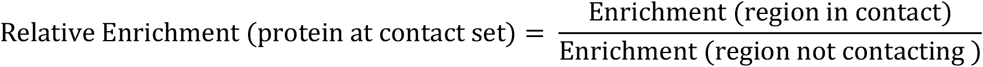

For Figs. 3B, 3E, 5C and 6B, the mean *relative enrichment* of the first minute of Jurkat T cell/target interaction was calculated. For Fig. 4B, *enrichment* of Lck in the segmented membranes was computed as above. In Fig. 4C, the mean *enrichment* of Lck averaged within the first minute of Jurkat T cell/target interaction was calculated.

### Statistics and Reproducibility

Graphpad Prism 9.3.0 was used to generate all graphs and to perform statistical analysis. Data distribution was assumed to be normal, but not formally tested. Data sets containing continuous data from different biological replicates (Fig. 1C, 1E, Supplementary Fig. 1B and Supplementary Fig. 2B) were presented as superplots (Lord et al., 2020). All statistical tests used are indicated in the figure legends. No statistical method was used to predetermine sample size and sample sizes are indicated in the figure legends. The experiments were not randomised. The Investigators were aware of the group allocation during experiments and outcome assessment. All schematics were generated with Affinity Designer. Microscopy images and Western Blots are shown as representative images.

## Data availability

All data supporting the findings of this study are available from the corresponding author on reasonable request.

## Acknowledgements

This project was supported by the Human Frontier Science Program early career grant for X.S. and F.B. Additional funding support for X.S. includes the American Cancer Society Research (Grant 135926), the NIGMS MIRA program (R35 GM138299), the Gabrielle’s Angel Foundation Medical Research Award and the Pershing Square Sohn Prize for Young Investigators in Cancer research. C.R. was supported by a PhD fellowship from SFB958, Deutsche Forschungsgemeinschaft (DFG). A.Z. is supported by the Deutsche Forschungsgemeinschaft – Project Number 278001972 – TRR 186. We thank Steffen Gottshalk for providing the HER2-CAR sequence employed in this study. We thank L. Lavis (Janelia Research Campus) for providing dyes for live-cell imaging. We thank Yvonne Weber (Freie Universität Berlin) for help with FACS. We thank Petia Adarska for help with imaging and for her valuable comments on the manuscript. We thank Steffen Erdle for cloning of the HER2-CAR-Halo plasmid. We thank Pia Hagenbach and Lucie Hortmann for valuable support in experiments.

## Author Contributions Statement

C.R., X.S. and F.B. conceived the project. C.R., G.C., E.F., H.A., K.D. X.S. and F.B. designed and performed experiments. A.Z. and C.R. and H.E. performed image analysis. C.R., G.C., E.F., H.A. and K.D. generated plasmids and knock-in cell lines. C.R. and F.B. wrote the manuscript with input from all authors.

