## Supplementary Figures 1-5 for "T cell protrusions enable fast, localised initiation of CAR signalling"

**Supplementary Information**

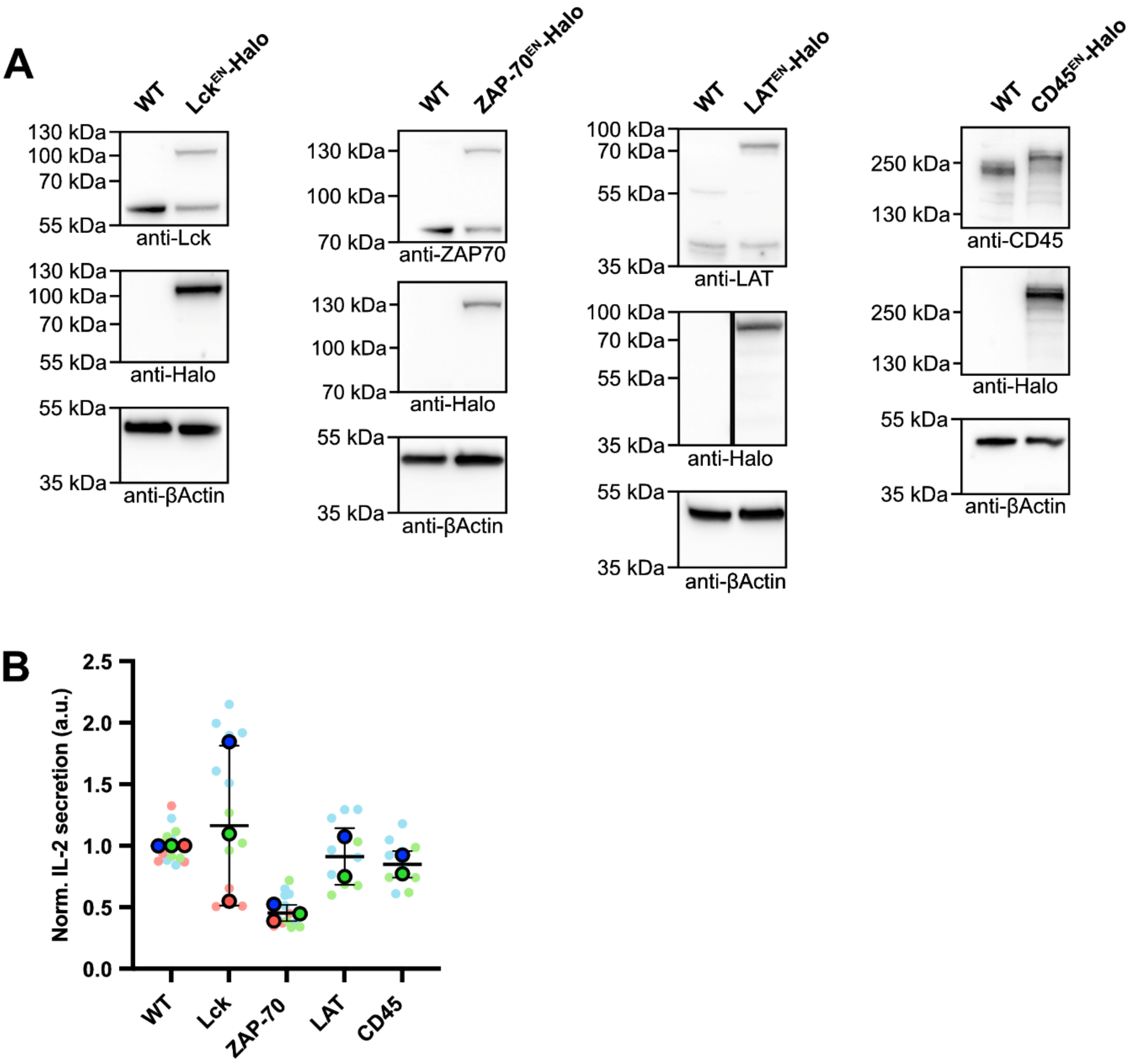


**Supplementary Figure 1. Validation of knock-in (KI) cell lines via Western Blot and functional IL-2 secretion assays. A)** Western blots of lysates of Jurkat T cell lines expressing endogenously Halo-tagged Lck, ZAP-70, LAT or CD45. Primary antibodies used for each immunoblot are shown below each crop. Full blots are provided in the source data file. All fusion proteins display the correct shift in molecular weight, corresponding to the molecular weight of the linker (short GS linker for Lck and LAT; long 70aa LAP linker for CD45 and ZAP-70), Halo and an epitope tag (2xalfa or 2xHA). For the CD45 KI all alleles appear to be edited as shown by a full shift of the band corresponding to WT CD45. For the Lck, ZAP-70 and LAT KI cell lines, only ~ 50% of alleles are edited. **B)** IL-2 secretion was assessed in supernatants of WT and KI Jurkat T cells cultured in dishes coated with OKT3 and CD28.2 antibodies for 24h. IL-2 secretion is normalised to the mean secretion from unmodified Jurkat T cells from the same day. Replicates are shown in different colours and each small dot represents the normalized concentration of IL-2 collected from the supernatant of one well. Graph shows mean values, s.d. error bars. P-values from paired t-tests are 0.7050 (WT/Lck), 0.050 (WT/ZAP-70), 0.6840 (WT/LAT) and 0.2970 (WT/CD45).

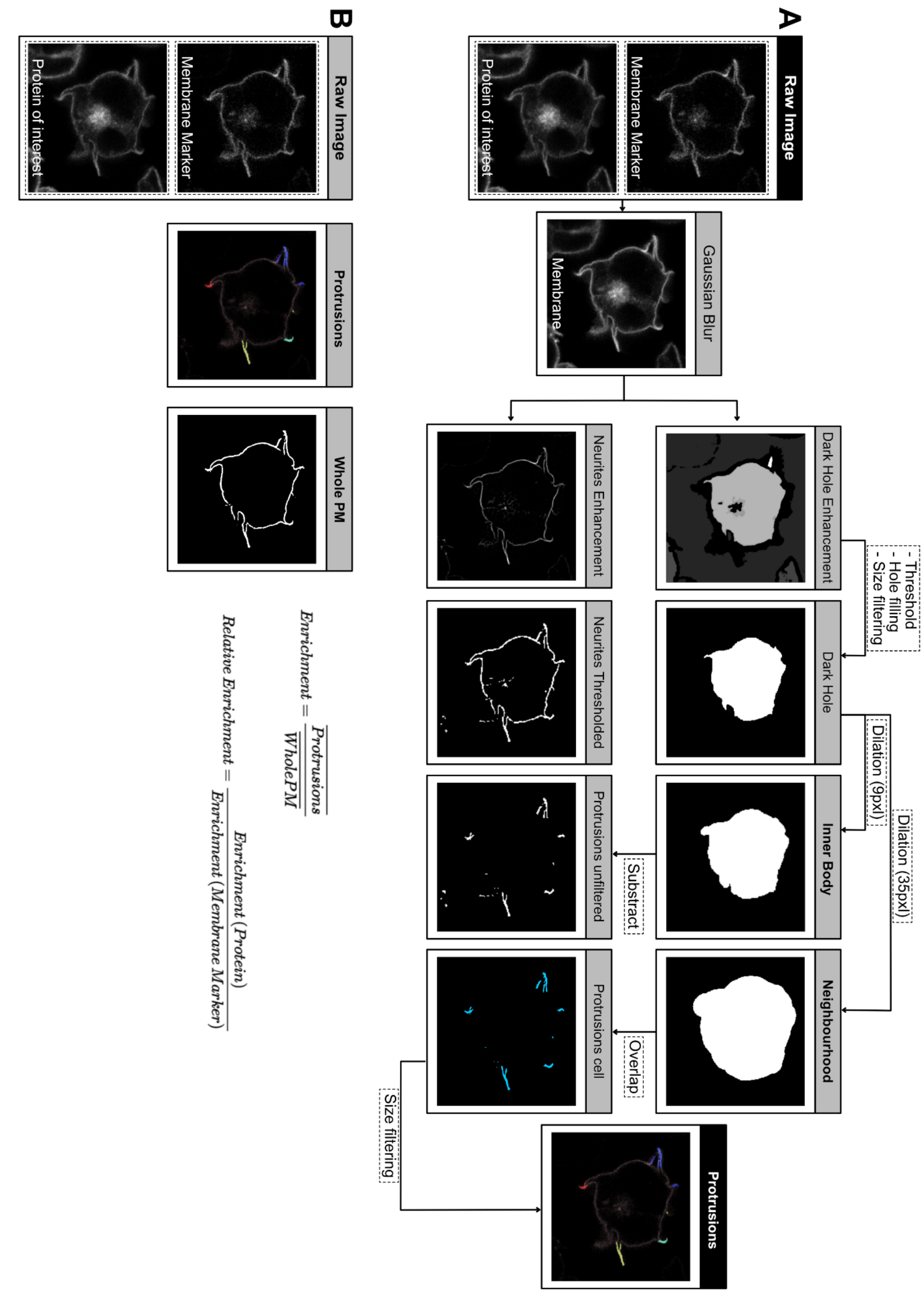


**Supplementary Figure 2. A)** Description of the protrusion segmentation pipeline. Two-colour images of proteins localizing to the plasma membrane of Jurkat T cells were used to segment protrusions using a custom CellProfiler 4 pipeline. First, the two membrane channels were averaged and gaussian blurred to generate a "membrane" image. (Upper row) For inner body detection, dark holes were enhanced, thresholded, size-filtered and filled. We then dilated the inner body 9 pixels, so it covered the main body membrane, and 35 pixels so we obtained the cell neighborhood. (Lower row) For membrane detection, neurite-like structures were enhanced and thresholded to obtain an unfiltered picture of the membrane. Unfiltered protrusions were obtained by subtracting the dilated inner body from the thresholded neurites. Protrusions outside the neighborhood were filtered out and size filtering was then applied for discarding single-pixel objects. **B)** Calculation of enrichment and relative enrichment of a protein of interest in a membrane set.

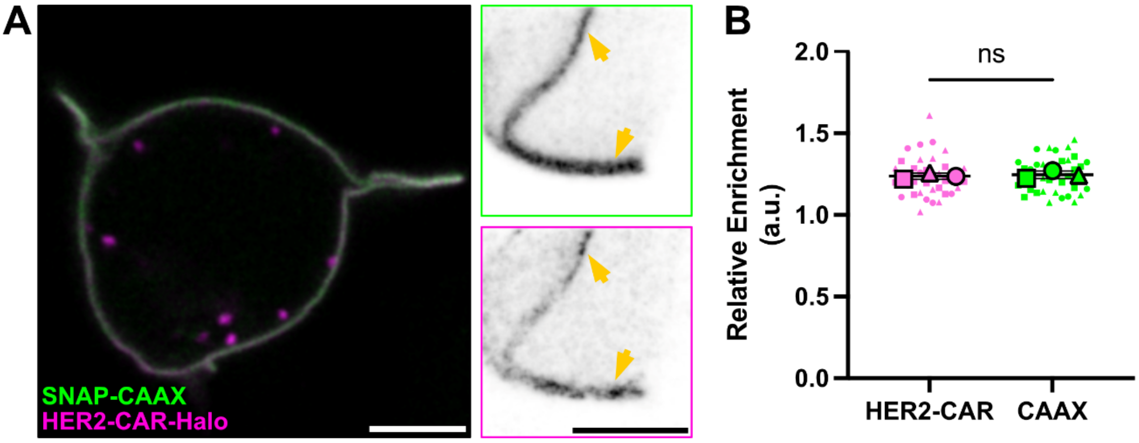


**Supplementary Figure 3. HER2-CAR shows no preferential localization to actin protrusions in resting conditions. A)** Live-cell confocal (magenta and green) and STED images (inverted greyscale) of a Jurkat T cell expressing HER2-CAR-Halo labelled with CA-$JFX_{650}$ and SNAP-CAAX labelled with BG-$JF_{571}$. Arrows highlight small HER2-CAR clusters localized either to the main body membrane or to an actin protrusion. **B)** Enrichment of HER2-CAR tagged with Halo in protrusions. In total, 37 cells from three independent experiments were analysed. Replicates are shown in different shapes, and each small dot represents a single cell. Graph shows mean values, standard deviation error bars. P-value of paired t-tests is 0.6116. CA = chloroalkane (HaloTag substrate), BG = benzylguanine (SNAP-tag substrate). Scale bars, 5 µm (confocal overview), 2 µm (STED images).

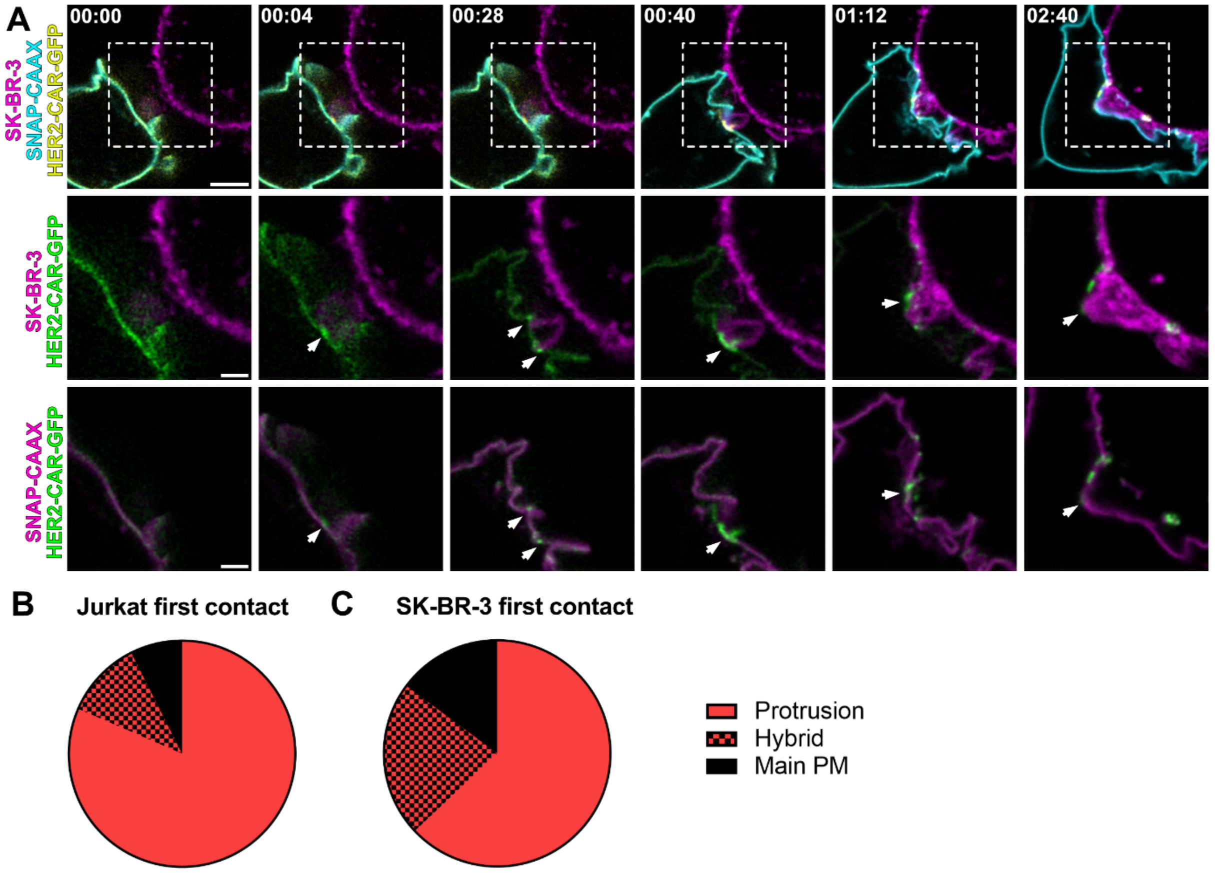


**Supplementary Figure 4. Protrusions from the target cell make contact with Jurkat T cells and lead to successful CAR engagement and clustering. A)** Time-lapse confocal imaging of a Jurkat T cell expressing HER2-CAR-GFP, SNAP-CAAX (labelled with BG-$JF_{571}$) interacting with a SK-BR-3 cell (labelled with CellMaskOrange). Dashed line describes the outline of the SK-BR-3 cell. Arrows highlight the cell-cell contact mediated by a protrusion emanating from the SK-BR-3. The contact leads to successful clustering of the HER2-CAR on the Jurkat T cell membrane. B) Number of Jurka T cell/SK-BR-3 first contacts mediated through Jurkat T cell protrusions, main body membrane or both at the same time. n = 29 from three independent experiments are represented. C) Number of Jurkat T cell/SK-BR-3 first contacts mediated through SK-BR-3 protrusions, main body membrane or both at the same time. n = 29 from three independent experiments are represented. BG = benzylguanine (SNAP-tag substrate). Scale bars, 5 µm (confocal overviews), 2 µm (crops).

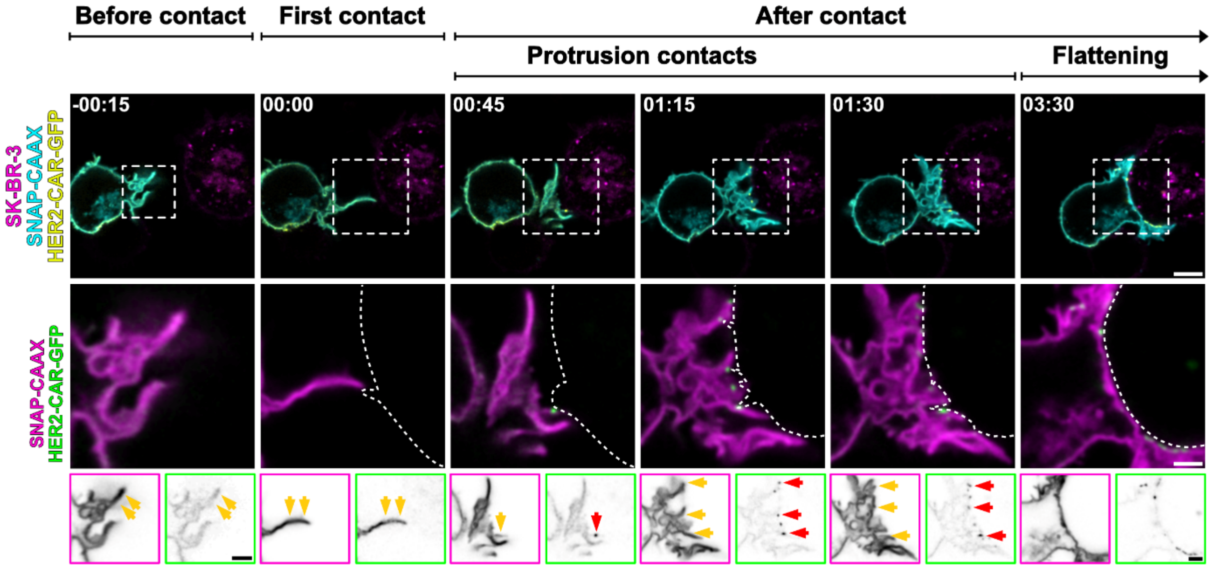


**Supplementary Figure 5. Early contact between a CD4+ primary T cells and the target cell is mediated by protrusions that trigger CAR clustering and activation.** Time-lapse confocal imaging of a CD4+ human T cell expressing HER2-CAR-GFP, SNAP-CAAX (labelled with BG-$JF_{571}$) interacting with a SK-BR-3 cell (labelled with CellMaskOrange). Dashed line describes the outline of the SK-BR-3 cell. HER2-CAR shows no preferential localization to protrusions prior to first contact with the target at t = 0s but is enriched in protrusions contacts. BG = benzylguanine (SNAP-tag substrate). Scale bars, 5 µm (confocal overviews), 2 µm (crops).
